# vsRNAfinder: a novel method for identifying high-confidence viral small RNAs from small RNA-Seq data

**DOI:** 10.1101/2022.05.30.493989

**Authors:** Zena Cai, Ping Fu, Ye Qiu, Aiping Wu, Gaihua Zhang, Yirong Wang, Taijiao Jiang, Xing-Yi Ge, Haizhen Zhu, Yousong Peng

**Affiliations:** Bioinformatics Center, College of Biology, Hunan Provincial Key Laboratory of Medical Virology, Hunan University, Changsha 410082, China; Institute of Systems Medicine, Chinese Academy of Medical Sciences & Peking Union Medical College, Beijing, China; Suzhou Institute of Systems Medicine, Suzhou, China; The National and Local Joint Engineering Laboratory of Animal Peptide Drug Development, College of Life Sciences, Hunan Normal University, Changsha 410081, China; Guangzhou Laboratory, Guangzhou, China

## Abstract

Virus-encoded small RNAs (vsRNA) have been reported to play an important role in viral infection. Unfortunately, there is still a lack of an effective method for vsRNA identification. Herein, we presented vsRNAfinder, a de novo method for identifying high-confidence vsRNAs from small RNA-Seq (sRNA-Seq) data based on peak calling and Poisson distribution and is public available at https://github.com/ZenaCai/vsRNAfinder. vsRNAfinder outperformed two widely-used methods namely miRDeep2 and ShortStack in identifying viral miRNAs with a significantly improved sensitivity. It can also be used to identify sRNAs in animals and plants with similar performance to miRDeep2 and ShortStack. The study would greatly facilitate effective identification of vsRNAs.

## Introduction

Small RNAs (sRNAs) are a kind of regulatory RNAs in cellular processes[1]. They are typically less than 200 nucleotides (nt) in length[1]. The sRNAs can be mainly divided into three classes including microRNA (miRNA) with a length of 18-24 nt, small interfering RNA (siRNA) with a length of 21-23 nt and piwi-interacting RNA (piRNA) with a length of 24-31 nt based on the mechanism of biogenesis and function[2]. By the mechanism of biogenesis, the miRNAs are derived from single-stranded RNA precursors with a hairpin structure which are firstly processed by Drosha in the nucleus and then processed by Dicer after being transported to the cytoplasm; the siRNAs are derived from double-stranded RNA precursors in the cytoplasm and are processed by Dicer; the piRNAs are derived from the fragmentation of single-stranded RNA precursors via a Dicer-independent mechanism[3-5]. By the mechanism of function, although all three kinds of sRNAs can target the mRNAs to regulate their expressions by incorporating sRNAs into the RNA-induced silencing complex (RISC), their function mechanisms differ a lot: with the help of AGO protein, the miRNAs can repress mRNAs translation by binding imperfectly to their mRNA targets, while the siRNAs can degrade mRNAs by binding perfectly to their mRNA targets; the piRNAs can degrade mRNAs by binding perfectly their mRNA targets with the help of PIWI protein[5, 6]. The sRNAs play a fundamental role in the maintenance of normal physiology and serve as potential biomarkers for early diagnosis and therapeutic targets for a wide spectrum of diseases, particularly the cancer[1]. For example, miR-130 can down-regulate the expression of miRNA-221 and miRNA-222 by targeting the MET, thus reducing TNF-related apoptosis-induced ligand (TRAIL) resistance in non-small cell lung cancer[7].

Numerous methods have been developed to identify sRNAs from sRNA-Seq data. They can be mainly divided into two kinds. The first kind of method is to identify sRNAs based on a known sRNA library, such as the sRNAbench[8], Manatee[9] and sRNAnalyzer[10]. This kind of method can be used for accurate quantification of known sRNA. However, they may not be suitable for organisms with only a few known sRNAs such as viruses. The other kind of method is to identify sRNAs *de novo* by firstly finding the precursor of sRNAs from assembly of reads and then detecting the mature sRNA, such as miRDeep2[11] and ShortStack[12]. Most methods mentioned above have been mainly used in the identification of sRNAs in animals and plants. The viruses have much smaller genomes and stronger coding ability compared to animals and plants. Thus, the distribution of vsRNAs is much denser than that of other organisms, which may lead to assembly of adjacent vsRNAs with overlaps into long contigs and missed detection of many vsRNAs by current methods.

Herein, we presented vsRNAfinder, a *de novo* method for the identification of high-confidence vsRNAs from sRNA-Seq data based on peak calling and Poisson distribution. The method can also be used in the identification of sRNAs in animals and plants. The work would greatly facilitate the identification of vsRNAs.

## Materials and methods

### Data collection

To evaluate the performance of computational methods in identifying viral miRNAs, a total of 196 known miRNAs in nine viruses were obtained from the miRBase database (Release 22.1, http://mirbase.org/) (Table S1). The viral-infection-related sRNA-Seq data of these viruses were obtained from the NCBI GEO[13] and SRA[14] databases. Only the sRNA-Seq data with the study type of “Expression profiling by high throughput sequencing” or “Non-coding RNA profiling by high throughput sequencing” in GEO database, and those with the strategy of “RNASeq” or “other” in SRA database were kept. Then, the sRNA-Seq data were further filtered manually. Only those of viral infections with the genome sequence of the virus used in the infection available were kept for further analysis. A total of 14 sRNA-Seq datasets for nine viruses were used in the study (Table S1).

To evaluate the performance of computational methods in identifying viral siRNAs and piRNAs, the experimentally-verified siRNAs in Sugarcane mosaic virus and piRNAs in Dengue viruses were collected from the studies of Xia et al.[15] and Wang et al.[16], respectively. The sRNA-Seq datasets of these two viruses were obtained from the NCBI GEO and SRA database as mentioned above (Table S2).

To demonstrate the usage of our method in the identification of sRNAs from other organisms, the known miRNAs in several widely-studied organisms including *Homo sapiens, Mus musculus, Drosophila melanogaster, Arabidopsis thaliana* and *Oryza sativa* were obtained from the database of miRBase. Some representative sRNA-Seq datasets of these organisms were also obtained from NCBI GEO and SRA databases (Table S3).

### The vsRNAfinder workflow

The vsRNAfinder workflow included four modules: Preprocessing, Identification, Filtering and annotation, and Quantification, which were described as follows.

#### Preprocessing module

Firstly, the adapter sequences are trimmed from the raw reads with cutadapt (version 2.6, parameter: ‘-m 15’)[17] (Figure 1). Then, the clean reads are mapped to the viral genome directly (the default), or the clean reads are firstly mapped to the host genome and then the unmapped reads are mapped to the viral genome, with bowtie (version 1.3.0, parameter: ‘-v 0 -m 2’)[18]. The resulted SAM files are converted to BAM files which are sorted by samtools (version 1.9)[19]. The sorted BAM files are further converted to BED files by bedtools (version 2.29.2)[20].

**Figure 1.**
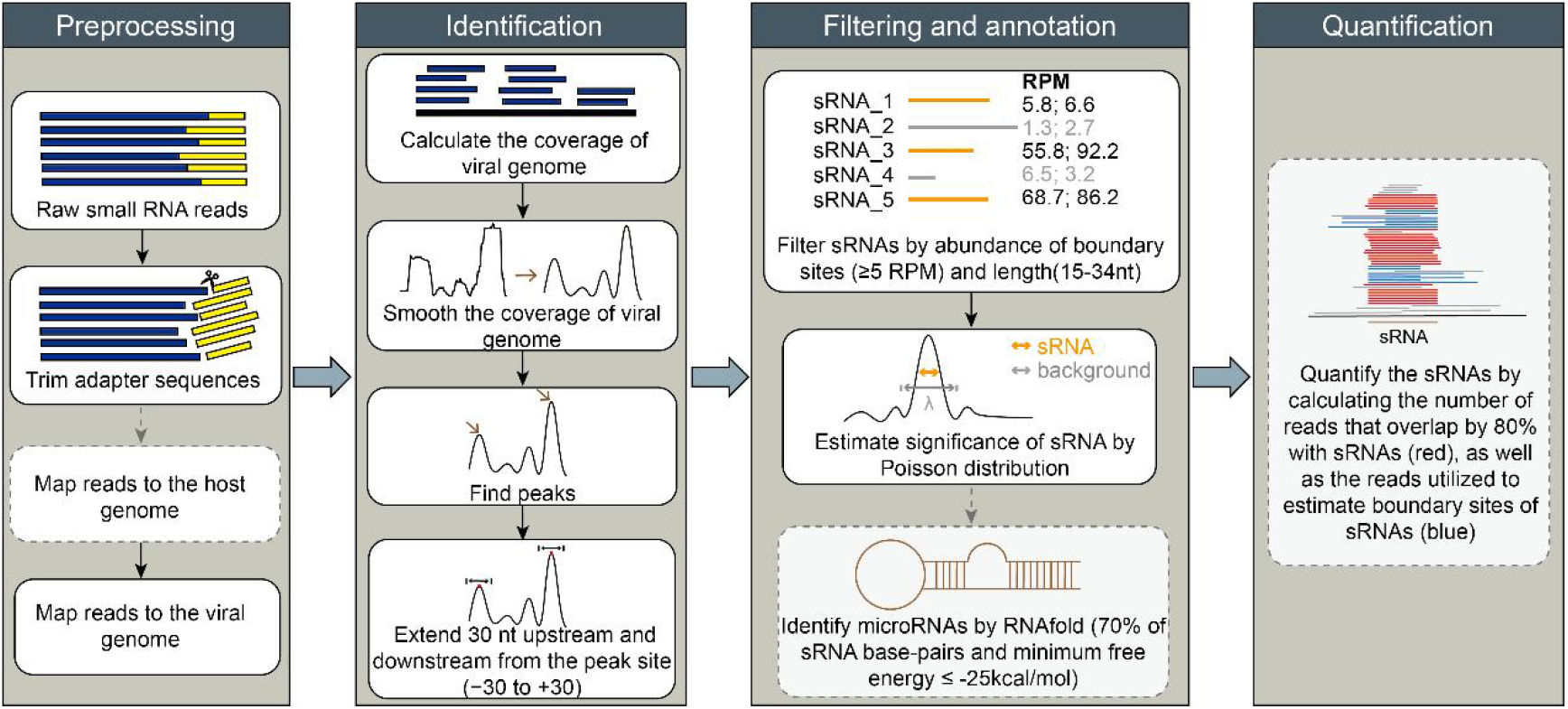
Overview of vsRNAfinder which consisted of four modules including Preprocessing, Identification, Filtering and annotation, and Quantification. Please see Materials and methods for details.

#### Identification module

The coverage of mapped reads on the viral genome is calculated by bedtools and then smoothed with the window-size of 11 nt. The peaks of the smoothed curve are identified using the function of *findPeaks* (“minpeakheight = 5”) in the R package pracma (version 2.3.3)[21]. The start and end site of the candidate sRNA is restricted to the upstream and downstream 30 nt of the peak, respectively. Then, the number of reads starting at each position in the upstream region of the peak, and that ending at each position in the downstream of the peak are calculated. The position with the highest number of reads in the upstream and that in the downstream are considered as the start and end site of the candidate sRNA, respectively.

#### Filtering and annotation module

To obtain the high-confidence sRNAs, the abundances of the start and end sites (defined as boundary sites) of the candidate sRNA are normalized using RPM (Reads Per Million) which is calculated by dividing the number of reads starting at the start site or ending at the end site by the total number of clean reads, and then multiplied by 10^6^. Only the candidate sRNAs with both of the boundary sites having at least 5 RPM and with lengths ranging from 15 to 34 nt are retained for further analysis.

Then, the Poisson distribution is used to evaluate the statistical significance of candidate sRNAs. Poisson distribution is a discrete probability distribution that expresses the probability of a given number of events occurring in a fixed interval of time, which is widely applied in NGS data analysis[22]. We hypothesized that the reads should be enriched in the region where the sRNA is expressed compared to the background region. Thus, the significance of candidate sRNAs could be analyzed by the Poisson distribution of which the parameter λ can be calculated as the average coverage of the mapped reads on the background region. For each candidate sRNA, the background region is defined as extending the upstream and downstream 10 nt of the candidate sRNA. Therefore, the λ is local and is calculated as follows:

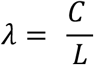

where C is the sum of the coverage of mapped reads on each position of the background region, and L is the length of the background region.

Based on the local λ, for a given position X in the candidate sRNA with *k* mapped reads covering the position, the p-value of the position is calculated as follows:

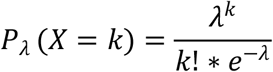

The p-value of the candidate sRNA is calculated as the 80% quantile of p-values in all positions of the candidate sRNA. The candidate sRNAs with a p-value smaller than 0.05 are considered as statistically significant and highly confident.

The miRNAs are then annotated from the high-confidence sRNAs in two steps. Firstly, the hypothesized precursor of miRNA (pre-miRNA) is obtained. Since the miRNA may be located in either 5’-arm or 3’-arm of the hypothesized pre-miRNA, both cases are considered. If the miRNA is supposed to be in the 5’-arm of the hypothesized pre-miRNA, the start and end positions of hypothesized pre-miRNAs are determined as follows according to Hackenberg et al.’s study[23]:

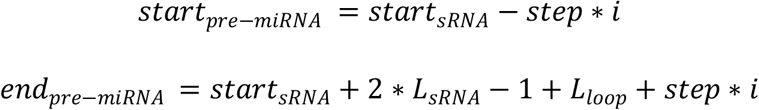

otherwise,

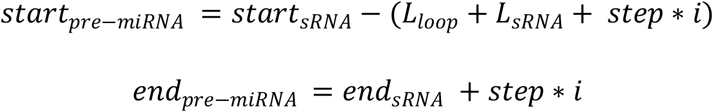

where *start*_*sRNA*_ and *end*_*sRNA*_ refer to the start and end position of the sRNA, respectively; *L*_*sRNA*_ refers to the length of sRNA; *L*_*loop*_ refers to the length of loop and is set to be 15; *step* and *i* are set to be 5 and 8 for animals and viruses, 7 and 10 for plants as plant pre-miRNA sequences are longer than those of animals and viruses, according to Hackenberg et al.’s study[23].

Then, the secondary structure of the hypothesized pre-miRNA is predicted by RNAfold (version 2.4.18)[24] with default parameters. The hypothesized pre-miRNA which has the minimum free energy of less than -25 kcal/mol, and in which at least 70% of positions in the corresponding sRNA form base pairs, is considered to be pre-miRNA having the hairpin structure, and is further processed into the mature miRNA.

#### Quantification module

The abundances of sRNAs are normalized using RPM. The total number of reads mapping to the sRNA is calculated as the sum of reads that overlap with the sRNA by 80% and those that are utilized to estimate boundary sites of the sRNA. The RPM of each sRNA is calculated by dividing the total number of reads mapping to the sRNA by the total number of clean reads in the sample, and then multiplied by 10^6^.

### Prediction of sRNAs with miRDeep2 and ShortStack

The miRDeep2 (version 0.1.2, https://github.com/rajewsky-lab/mirdeep2)[11] and ShortStack (version 3.8.5, https://github.com/MikeAxtell/ShortStack)[12] were used for prediction of sRNAs with default parameters.

### Evaluating the performance of computational methods in identifying sRNAs

The known sRNAs (mentioned in Data collection) including miRNAs, siRNAs and piRNAs were used to evaluate the performance of computational methods in identifying sRNAs. For a given organism, the identified sRNAs whose sequences match perfectly to those of known sRNAs are considered as true positives, otherwise false positives; the known sRNAs which are missed by the method are considered as false negatives. Three metrics including Precision, Recall and F1-score were used to evaluate the performance of computational methods in identifying sRNAs, which are calculated as follows:

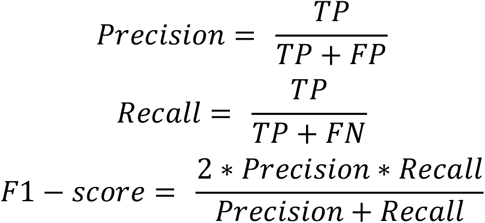

while TP is the number of true positives, FP is the number of false positives, FN is the number of false negatives.

### Statistical analysis

The Wilcoxon signed-rank test was used to compare the F1-score between the vsRNAfinder and the other methods. A p-value less than 0.05 was considered as statistically significant. All statistical analyses were conducted in Python (version 3.8).

## Results

### Overview of vsRNAfinder

The vsRNAfinder consisted of four modules including Preprocessing, Identification, Filtering and annotation, and Quantification (Figure 1). The Preprocessing module processed raw reads to clean reads by trimming adapters. Then, the clean reads were firstly mapped to the host genome and then the unmapped reads were mapped to the viral genome, or the clean reads were mapped to the viral genome directly (the default) since the host reference genome was not always available.

The Identification module identified the candidate sRNAs with four steps: firstly, the coverage of mapped reads across the viral genome was calculated; secondly, the coverage was smoothed to reduce the noise of the peak detection and enhance the peak signal; thirdly, the peak was detected which had a higher expression than its adjacent sites; finally, the start and end sites of the candidate sRNA were determined at the upstream and downstream of the peak identified, respectively (see Materials and methods).

The Filtering and annotation module was designed for obtaining high-confidence sRNAs and annotating miRNAs. The candidate sRNAs identified above were firstly filtered by the abundance of the start and end sites (defined as boundary sites) of sRNAs. Only the candidate sRNAs with both boundary sites having at least 5 RPM were kept. Then, they were further filtered by the length. Only the candidate sRNAs with 15-34 nt were kept which covered the length of most miRNAs, siRNAs and piRNAs[2]. Then, the statistical significance of candidate sRNAs was obtained based on the Poisson distribution. The remaining candidate sRNAs with a p-value smaller than 0.05 were considered as statistically significant and highly confident. Finally, the RNAfold[24] was used to annotate miRNAs from the high-confidence sRNAs (see Materials and methods).

The Quantification module quantified the abundance of sRNAs by RPM. The total number of reads mapping to the sRNA was calculated as the sum of reads that overlapped with the sRNA by 80% and those that were utilized to estimate boundary sites of the sRNA (see Materials and methods).

### Performance analysis of vsRNAfinder in identifying viral miRNAs

To evaluate the performance of vsRNAfinder in identifying vsRNAs, vsRNAfinder was used to identify viral miRNAs since the miRNAs are the most studied vsRNAs. Nine viruses which had a range of 6-44 miRNAs reported in miRBase were used in the evaluation (Figure 2). For comparison, two *de novo* methods, namely the miRDeep2[11] and ShortStack[12] which are widely used in identifying sRNAs, were selected in identifying viral miRNAs in nine viruses.

**Figure 2.**
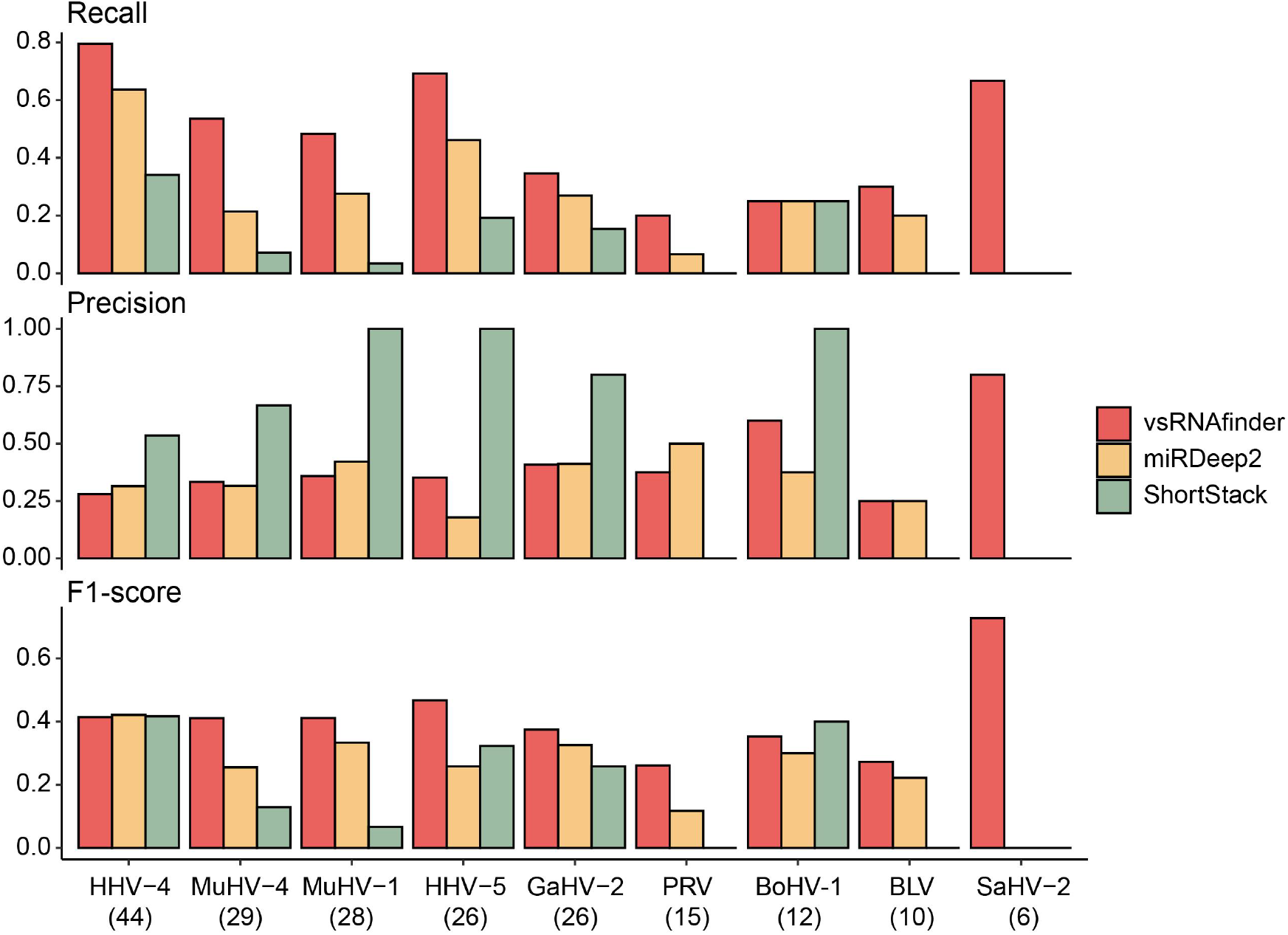
Performance analysis of vsRNAfinder, miRDeep2 and ShortStack in identifying viral miRNAs. Comparison of Recall, Precision and F1-score of vsRNAfinder with miRDeep2 and ShortStack in identifying viral miRNAs in viruses of Human gammaherpesvirus 4 (HHV-4), Murid gammaherpesvirus 4 (MuHV-4), Murid betaherpesvirus 1 (MuHV-1), Human betaherpesvirus 5 (HHV-5), Gallid alphaherpesvirus 2 (GaHV-2), Suid alphaherpesvirus 1 (PRV), Bovine alphaherpesvirus 1 (BoHV-1), Bovine leukemia virus (BLV) and Saimiriine gammaherpesvirus 2 (SaHV-2). The numbers in parentheses under the viral name referred to the number of miRNAs reported in miRBase.

As shown in Figure 2, vsRNAfinder achieved the highest Recall ranging from 0.2 to 0.8 in all viruses, followed by miRDeep2 (0-0.64) and ShortStack (0-0.34). Taking the MuHV-4 as an example, vsRNAfinder had a Recall of 0.54, while miRDeep2 and ShortStack had a Recall of 0.21 and 0.07, respectively. In terms of Precision, the ShortStack performed best with Precision higher than 0.5 in all viruses except three viruses for which no miRNAs were identified by ShortStack. The vsRNAfinder had Precisions ranging from 0.25-0.8 which were similar to those of miRDeep2. Overall, when considering both the Recall and Precision using the F1-score, the vsRNAfinder achieved the best F1-scores in all viruses except HHV-4 and BoHV-1. It had a median F1-score of 0.41 which was significantly larger than that of miRDeep2 (0.26, p-value = 0.0098) and ShortStack (0.13, p-value = 0.0039).

When comparing the number of viral miRNAs identified by three methods, we found that most viral miRNAs identified by miRDeep2 and ShortStack were also identified by vsRNAfinder (Table S4). For example, 19 and 3 viral miRNAs were identified by miRDeep2 and ShortStack, respectively, in the MuHV-4, among which 14 and 3 were also identified by vsRNAfinder. Moreover, vsRNAfinder identified more novel viral miRNAs than the other two methods in most viruses (Table S4).

### Identification of other classes of sRNAs with vsRNAfinder

Besides viral miRNAs, viruses can also express siRNAs and piRNAs during viral infections[15, 16]. Although vsRNAfinder could not annotate siRNAs and piRNAs from vsRNAs, it could identify them from sRNA-Seq data. To evaluate the performance of vsRNAfinder in the identification of other vsRNA classes, 11 experimentally-verified siRNAs in Sugarcane mosaic virus (SCMV)[15] and 6 experimentally-verified piRNAs in Dengue virus (DENV)[16] were collected from literature. All SCMV siRNAs and four of six DENV piRNAs were identified correctly by vsRNAfinder (Table S5). The remaining two DENV piRNAs were missed by vsRNAfinder due to low abundance. For comparison, ShortStack only identified correctly one SCMV siRNA and two DENV piRNAs (Table S5), while miRDeep2 can only be used to identify the miRNA.

### Performance analysis of vsRNAfinder in identifying sRNAs in plants and animals

To investigate the ability of vsRNAfinder in identifying sRNAs in other organisms besides viruses, the vsRNAfinder was used to identify sRNAs in *Homo sapiens* and four model organisms including *Mus musculus, Drosophila melanogaster, Arabidopsis thaliana* and *Oryza sativa*. The ability of vsRNAfinder was evaluated based on its performance in identifying miRNAs in these organisms. For comparison, miRDeep2 and ShortStack were also used to identify miRNAs in these organisms. As shown in Table 1, vsRNAfinder had Recalls ranging from 0.06 to 0.13 in humans and two animals which were a little lower than those of miRDeep2, while it had Recalls of 0.05 to 0.11 in two plants which were a little higher than those of miRDeep2. When compared to ShortStack, vsRNAfinder consistently had higher Recalls in five organisms. In terms of Precision, the ShortStack had the highest Precision among the three methods in five organisms. Although the Precision of vsRNAfinder was lower than that of ShortStack, it was consistently higher than that of miRDeep2 in five organisms. Overall, in terms of F1-score which combined both the Recall and Precision, the vsRNAfinder had an average F1-score of 0.122 which was similar to that of miRDeep2 (0.116) and was slightly higher than that of ShortStack (0.098).

**Table 1.**
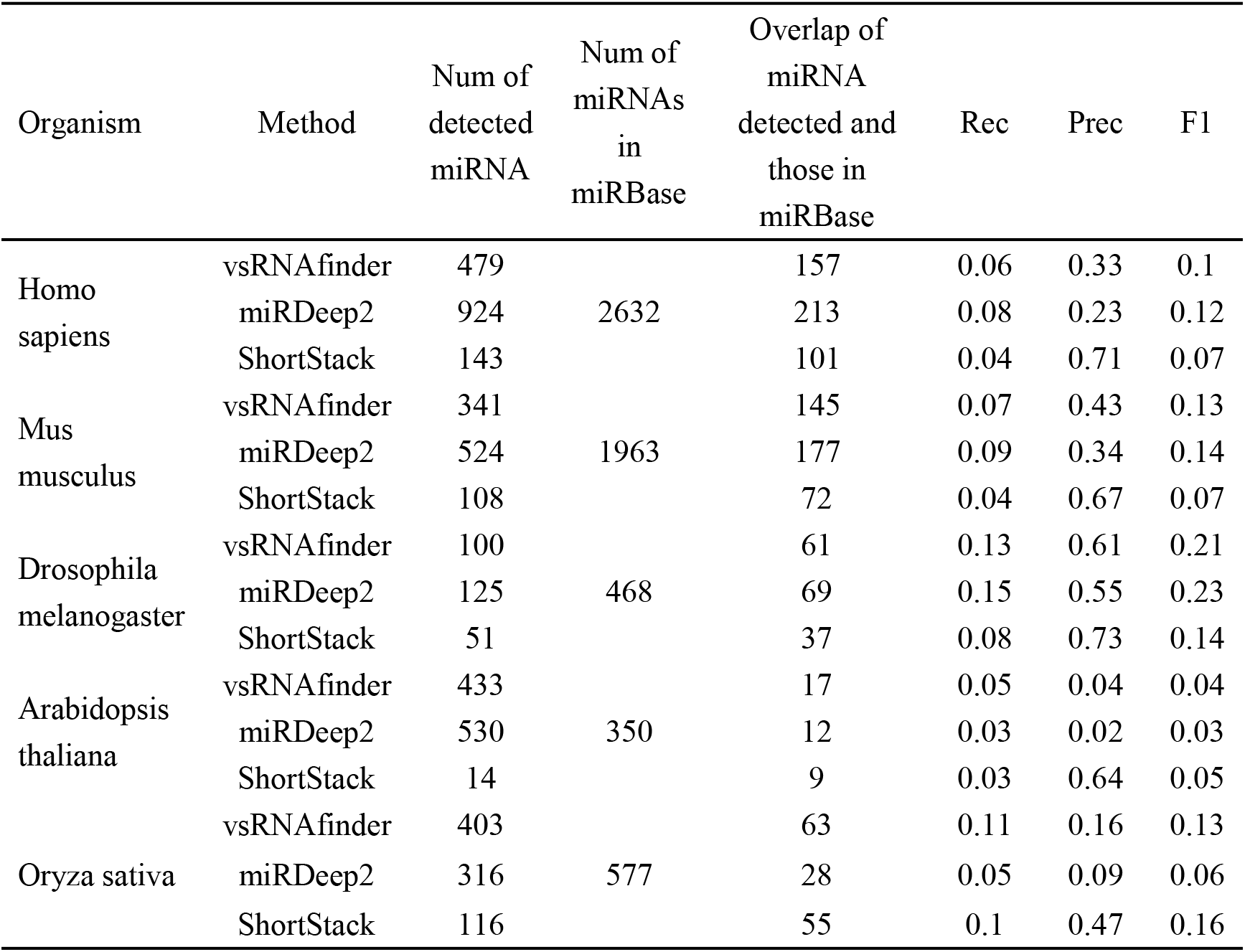
Performance comparison of three computational methods in identifying miRNAs in five organisms. Rec, Recall; Prec, Precision; F1, F1-score.

| Organism | Method | Num of detected miRNA | Num of miRNAs in miRBase | Overlap of miRNA detected and those in miRBase | Rec | Prec | F1 |
| --- | --- | --- | --- | --- | --- | --- | --- |
| Homo sapiens | vsRNAfinder | 479 | 2632 | 157 | 0.06 | 0.33 | 0.1 |
|  | miRDeep2 | 924 |  | 213 | 0.08 | 0.23 | 0.12 |
|  | ShortStack | 143 |  | 101 | 0.04 | 0.71 | 0.07 |
| Mus musculus | vsRNAfinder | 341 | 1963 | 145 | 0.07 | 0.43 | 0.13 |
|  | miRDeep2 | 524 |  | 177 | 0.09 | 0.34 | 0.14 |
|  | ShortStack | 108 |  | 72 | 0.04 | 0.67 | 0.07 |
| Drosophila melanogaster | vsRNAfinder | 100 | 468 | 61 | 0.13 | 0.61 | 0.21 |
|  | miRDeep2 | 125 |  | 69 | 0.15 | 0.55 | 0.23 |
|  | ShortStack | 51 |  | 37 | 0.08 | 0.73 | 0.14 |
| Arabidopsis thaliana | vsRNAfinder | 433 | 350 | 17 | 0.05 | 0.04 | 0.04 |
|  | miRDeep2 | 530 |  | 12 | 0.03 | 0.02 | 0.03 |
|  | ShortStack | 14 |  | 9 | 0.03 | 0.64 | 0.05 |
| Oryza sativa | vsRNAfinder | 403 | 577 | 63 | 0.11 | 0.16 | 0.13 |
|  | miRDeep2 | 316 |  | 28 | 0.05 | 0.09 | 0.06 |
|  | ShortStack | 116 |  | 55 | 0.1 | 0.47 | 0.16 |

## Discussion

Numerous methods have been developed to identify sRNAs from sRNA-Seq data. However, there is still a lack of an effective method for sRNA identification in viruses. The *de novo* methods such as miRDeep2 and ShortStack identified sRNAs based on reads assembly, which may merge reads from multiple adjacent sRNAs with overlaps. This became worse in viruses on whose genome the sRNAs were more densely distributed than those in other organisms. Hence, many vsRNAs may be missed by these methods. The vsRNAfinder presented in the study is also a *de novo* method and outperformed miRDeep2 and ShortStack in capturing viral miRNAs with improved Recalls. This may be explained by the following two reasons. Firstly, smoothing the coverage of mapped reads on the viral genome before the peak calling could reduce the noise of the peak detection and enhance the peak signal. Secondly, filtering sRNAs by abundance and evaluating the statistical significance of vsRNAs based on the Poisson distribution could help identify high-confidence vsRNAs. Therefore, it is suggested to put vsRNAfinder on priority in the identification of vsRNAs.

Most methods have been developed for the identification of specific sRNA class such as miRNA, or the identification of sRNAs in a specific organism such as humans[11, 23]. Compared to previous methods, vsRNAfinder could be used to identify multiple classes of sRNAs including miRNA, piRNA and siRNA. Besides viruses, vsRNAfinder also achieved similar performances to current methods in the identification of sRNAs across several animal and plant organisms. This suggested that vsRNAfinder could be taken as a universal method for the identification of sRNAs, especially for the vsRNAs.

There are some limitations to the study. Firstly, only miRNAs were annotated in vsRNAfinder as there were few effective methods of annotating other vsRNA types such as piRNA and siRNA. For example, viral piRNAs were reported to lack the characteristic ping-pong signature that is common in piRNAs of other organisms, which makes it difficult to distinguish between viral piRNA and siRNA[25, 26]. More computational methods are needed to annotate vsRNAs. Secondly, the reads that mapped to more than two locations of the genome were excluded to reduce false positives in vsRNAfinder, which may lower the abundance of vsRNAs. Fortunately, only 0.007% of reads were mapped to more than two locations in the viral genomes as the viral genomes generally have a low level of sequence repeating.

## Conclusion

Overall, this study developed a state-of-the-art method named vsRNAfinder which can not only be used for identifying high-confidence sRNAs in viruses with better performances than current methods, but also be used for sRNA identification in plants and animals.

## Supporting information

Supplementary materals

## Availability of data and materials

All data used in the study are available in supplementary materials. The vsRNAfinder is available at https://github.com/ZenaCai/vsRNAfinder.

## Authors’ contributions

Zena Cai and Ping Fu contributed equally to this work. Zena Cai developed the method, analyzed the data and wrote the draft; Ping Fu analyzed the data; Ye Qiu, Aiping Wu, Gaihua Zhang, Yirong Wang, Taijiao Jiang, Xing-Yi Ge, Haizhen Zhu and Yousong Peng supervised the project, reviewed and revised the manuscript; Yousong Peng designed the project, wrote and revised the manuscript.

## Acknowledgements

We thank people in the PengLab for helpful discussions on the manuscript.

