## Supplementary materals for "vsRNAfinder: a novel method for identifying high-confidence viral small RNAs from small RNA-Seq data"

### Supplementary Tables

**Table S1.** The viral-infection-related sRNA-Seq datasets used in evaluating the ability of computational methods in identifying viral miRNAs in this study.

| Project | Viral species | Abbreviation | Accession number of viral genome |
|---|---|---|---|
| PRJNA181011 | Bovine leukemia virus | BLV | NC_001414.1 |
| PRJNA115945 | Bovine alphaherpesvirus 1 | BoHV-1 | AJ004801.1 |
| PRJNA510993 | Gallid alphaherpesvirus 2 | GaHV-2 | NC_002229.3 |
| PRJDB3333 | Human gammaherpesvirus 4 | HHV-4 | V01555.2 |
| PRJNA177675 | Human gammaherpesvirus 4 | HHV-4 | V01555.2 |
| PRJNA274950 | Human gammaherpesvirus 4 | HHV-4 | NC_007605.1 |
| PRJNA486534 | Human gammaherpesvirus 4 | HHV-4 | NC_009334.1 |
| PRJNA148583 | Human betaherpesvirus 5 | HHV-5 | FJ616285.1 |
| PRJNA269099 | Human betaherpesvirus 5 | HHV-5 | FJ616285.1 |
| PRJNA157095 | Murid betaherpesvirus 1 | MuHV-1 | GU305914.1 |
| PRJNA127943 | Murid gammaherpesvirus 4 | MuHV-4 | NC_001826.2 |
| PRJNA153641 | Murid gammaherpesvirus 4 | MuHV-4 | NC_001826.2 |
| PRJEB5032 | Suid alphaherpesvirus 1 | PRV | NC_006151.1 |
| PRJNA292108 | Saimiriine gammaherpesvirus 2 | SaHV-2 | NC_001350.1 |

**Table S2.** The viral-infection-related sRNA-Seq datasets used in evaluating the ability of computational methods in identifying viral siRNA and piRNA in this study.

| Project | Viral species | Abbreviation | Accession number of viral genome |
|---|---|---|---|
| PRJNA352783 | Sugarcane mosaic virus | SCMV | AY042184.1 |
| PRJNA530798 | Sugarcane mosaic virus | SCMV | AY042184.1 |
| PRJNA236401 | Dengue virus | DENV | KM204118.1 |
| PRJNA266286 | Dengue virus | DENV | KM204118.1 |

**Table S3.** The sRNA-Seq datasets of organisms in animals and plants used in this study. For the datasets of *Homo sapiens* and *Arabidopsis thaliana*, only samples from the same stage (first trimester) or the same tissue (preglobular embryos) were used in the analysis as lots of samples from different stages or tissues were available.

| Accession number of dataset in GEO | Host species | Cell/Tissue | Accession number of samples |
|---|---|---|---|
| GSE164178 | Homo sapiens | Placenta | SRR13347521 … SRR13347525 |
| GSE97622 | Mus musculus | Bone Marrow derived macrophages | SRR5443053; SRR5443054 |
| GSE57687 | Drosophila melanogaster | Brain | SRR1287660; SRR1287661 |
| GSE132066 | Arabidopsis thaliana | Preglobular embryos | SRR9179035 … SRR9179037 |
| GSE136142 | Oryza sativa | Mesocotyl | SRR10010317 … SRR10010322 |

**Table S4.** Comparison of viral miRNAs identified by vsRNAfinder, miRDeep2 and ShortStack in nine viruses which had miRNAs reported in miRBase. "V", vsRNAfinder; "M", miRDeep2; "S", ShortStack.

| Virus | Number of miRNAs detected by the method | | | Number of novel miRNAs (not in miRBase) detected by the method | | | Proportion of miRNAs which were also detected by vsRNAfinder | |
|---|---|---|---|---|---|---|---|---|
| | V | M | S | V | M | S | M | S |
| BoHV-1 | 5 | 8 | 3 | 2 | 5 | 0 | 0.25 | 1.00 |
| BLV | 12 | 8 | 2 | 9 | 6 | 2 | 0.50 | 1.00 |
| GaHV-2 | 22 | 17 | 5 | 13 | 10 | 1 | 0.76 | 1.00 |
| HHV-4 | 125 | 89 | 28 | 90 | 61 | 13 | 0.71 | 0.93 |
| HHV-5 | 51 | 67 | 5 | 33 | 55 | 0 | 0.30 | 1.00 |
| MuHV-1 | 39 | 19 | 1 | 25 | 11 | 0 | 0.68 | 1.00 |
| MuHV-4 | 45 | 19 | 3 | 30 | 13 | 1 | 0.74 | 1.00 |
| PRV | 8 | 2 | 0 | 5 | 1 | 0 | 1.00 | 0.00 |
| SaHV-2 | 5 | 0 | 0 | 1 | 0 | 0 | 0.00 | 0.00 |

**Table S5.** The performance of vsRNAfinder in identification of virus siRNA and piRNA, and its comparison to ShortStack. [#]IDs of the vsRNA detected by vsRNAfinder. Please see Table S6 for detailed information of the vsRNA. [*] N, not identified by vsRNAfinder or ShortStack; Y, identified by ShortStack.

| sRNA type and viral species | Reference | Experimentally validated vsRNA | | | | | Whether detected by the method | |
|---|---|---|---|---|---|---|---|---|
| | | ID | Start | End | Strand | Sequences | vsRNAfinder[#] | Short Stack |
| siRNA/ SCMV | PMID: 24819114 | vsiRNA1 | Not provided | | + | CUCUUGGACUCACGUGACAUAC | SCMV_sRNA_239 | N[*] |
| | | | | | + | UCUUGGACUCACGUGACAUACA | | |
| | | vsiRNA2 | | | + | CACGGUGAAUGCAGAAGAACU | SCMV_sRNA_443 | N |
| | | | | | + | ACGGUGAAUGCAGAAGAACUAC | | |
| | | | | | + | CGGUGAAUGCAGAAGAACUAC | | |
| | | vsiRNA3 | | | + | AGGAUGGCACUGUUAGAAAUCC | SCMV_sRNA_547 | N |
| | | | | | + | AGGAUGGCACUGUUAGAAAUC | | |
| | | | | | + | GGAUGGCACUGUUAGAAAUCC | | |
| | | vsiRNA4 | | | + | AGAAAAAGCGUAGAUUUAGGC | SCMV_sRNA_297 | N |
| | | vsiRNA5 | | | + | AAGAGUUGGAAGGAACAAGCC | SCMV_sRNA_331 | N |
| | | | | | + | AGAGUUGGAAGGAACAAGCCU | | |
| | | vsiRNA6 | | | + | ACGGAGAUUUCUGGAAACACU | SCMV_sRNA_372 | N |
| | | vsiRNA7 | | | - | GUUGAGAGAGAAGAAUCAGAGA | SCMV_sRNA_451 | N |
| | | vsiRNA8 | | | - | AUCCCUGAUCAAUUUCGUGACG | SCMV_sRNA_62 | Y |
| | | vsiRNA9 | | | - | AAUAGGAUUUCUAACAGUGCC | SCMV_sRNA_71 | N |
| | | vsiRNA10 | | | - | GUAUUAACUUUGAUGAGCGCCU | SCMV_sRNA_6 | N |
| | | | | | - | GUAUUAACUUU | | |

| | | | | | | | | |
|---|---|---|---|---|---|---|---|---|
| | | | | | | GAUGAGCGCC | | |
| | | vsiRNA11 | | | - | AGUAUUAACUUUGAUGAGCGCC | SCMV_sRNA_19 | N |
| | | | | | - | CCAUAGCAACUUGUAGGGCUG | | |
| | | | | | - | UUCCAUAGCAACUUGUAGGGCU | | |
| | | | | | - | UCCAUAGCAACUUGUAGGGCU | | |
| | | | | | - | UUCCAUAGCAACUUGUAGGGC | | |
| | | vpiRNA1 | 961 | 982 | + | AGAGACUUUGUAGAAGGGGUUU | DENV_sRNA_2 | N |
| | | | 961 | 981 | + | AGAGACUUUGUAGAAGGGGUU | | N |
| | | vpiRNA2 | 1105 | 1125 | + | AGGAAGUACUGUAUAGAGGCA | N | N |
| | | vpiRNA3 | 4566 | 4587 | + | GGCUGAACUGGAAGAUGGAGCC | DENV_sRNA_5 | N |
| | | vpiRNA4 | 5719 | 5738 | + | AAUGAUUGGACUUCGUGGU | DENV_sRNA_22 | N |
| piRNA/ DENV | PMID: 29690553 | vpiRNA5 | 8037 | 8057 | + | AGAAGCAGGACGAACACUCAG | N | Y* |
| | | vpiRNA6 | 8331 | 8355 | + | AGAUGUAGACCUCGGAAGCGGAACC | DENV_sRNA_1;DENV_sRNA_30 | Y |
| | | | 8331 | 8351 | + | AGAUGUAGACCUCGGAAGCGG | | |
| | | | 8333 | 8353 | + | AUGUAGACCUCGGAAGCGGAA | | |
| | | | 8335 | 8355 | + | GUAGACCUCGGAAGCGGAACC | | |
| | | | 8335 | 8356 | + | GUAGACCUCGGAAGCGGAACCC | | |
